# ArchaeaHQ: A Curated Reference Database of Archaeal Genomes

**DOI:** 10.64898/2026.06.02.729493

**Authors:** Dmitry Bespiatykh, Pedro Leão

## Abstract

Archaea have proven to be major players in biogeochemical cycles across diverse ecosystems, yet we still see an underrepresentation of archaeal genomes in the datasets used by popular computational biology tools. Here we present ArchaeaHQ, a quality-controlled, systematically curated reference database of 21,644 archaeal genomes compiled initially from 35,993 assemblies from all four archaeal kingdoms retrieved from NCBI: Methanobacteriati (Euryarchaeota), Thermoproteati (TACK), Nanobdellati (DPANN), and Promethearchaeati (Asgard). All genomes in the database passed standardized quality control, requiring ≥70% completeness and ≤10% contamination. A total of 44.2% of genomes in ArchaeaHQ achieved ≥90% completeness, while 93.1% exhibited ≤5% contamination. ArchaeaHQ comprises 16,199 metagenome-assembled genomes (MAGs; 74.8%) and 5,445 isolate genomes (25.2%). Approximately 75% of MAGs are assigned to 17 ecologically meaningful categories based on sampling origin, and around 65% of genomes include geographic metadata. ArchaeaHQ is available at https://doi.org/10.6084/m9.figshare.32266599 and provides an analysis-ready reference set for metagenomic classification, biogeochemical and ecological studies, comparative genomics, and development of archaeal-specific bioinformatic tools.

**Impact Statement:** Archaea are key drivers of the global carbon, nitrogen and methane cycles, yet their genomes remain underrepresented and inconsistently curated in the public databases that power modern computational biology tools. We present ArchaeaHQ, a quality-controlled, systematically curated reference set of 21,644 archaeal genomes spanning all four archaeal kingdoms, each passing standardized completeness and contamination thresholds and enriched with environmental and geographic metadata. By providing an analysis-ready, downloadable resource compatible with standard pipelines, ArchaeaHQ fits the gap between taxonomy-focused frameworks and unfiltered genome archives supporting metagenomic classification, biogeochemical and ecological studies, comparative genomics, and the development of archaeal-specific bioinformatic tools.

## Introduction

From the production of methane in water bodies to the oxidation of ammonia in the ocean, Archaea drive biogeochemical processes that have been shaping life on Earth from its inception to modern days. Today, Methanogenic archaea are responsible for the production of approximately 1 Gt of methane per year [1]. On the other side, Anaerobic methane-oxidizing archaea (ANME) operate in syntrophy with sulphate-reducing bacteria at the sediment-water interface consuming methane ascending from marine sediments, working as a natural filter reducing methane emissions [2, 3]. As key contributors to the global methane cycle, the metabolism of archaea within the kingdom Methanobacteriati (Euryarchaeaota) must be a main topic of discussion when we investigate solutions for climate crises. Ammonia-oxidizing archaea are representatives from the kingdom Thermoproteati (TACK) and are known for catalyzing the conversion of ammonia to nitrite, a key step in nitrification and thereby the coupling carbon and nitrogen cycles at global scales [4, 5].

In the last 10 years the metagenomic revolution revealed that the cultivated fraction of archaeal diversity represents a minor, phylogenetically skewed, sample of what exists in nature [6, 7]. Among other discoveries, metagenomics introduced us to two new archaeal kingdoms: Nanobdellati (DPANN) and Promethearchaeati (Asgard). DPANN archaea are ultra-small archaea with extremely reduced genomes that appear to live in synergy (as episymbionts or parasites) with other archaeal groups [8]. While Asgard archaea represent the closest known archaeal relatives of eukaryotes [9, 10] their study has continued to refine eukaryogenesis models driving us closer to uncover the origin of complex life on Earth [9, 11].

The impact of this revolution is reflected in public genome online resources. For example, in NCBI GenBank among 35,993 available archaeal genome assemblies as of February 2025, more than 70% of them are metagenome-assembled genomes (MAGs) recovered from environmental samples rather than cultivated isolates. Yet the addition of assemblies into public archives proceeds without systematic quality filtering, metadata standardization, or redundancy control, meaning that raw public archives are simultaneously a valuable scientific resource, and a risky analytical mixture of high-quality genomes, fragmented or contaminated assemblies, and redundant sequences with inconsistent or absent environmental data. Several genomic resources address some of these issues, but none were designed to serve as a dedicated, analysis-ready high-quality archaeal genome reference set.

NCBI GenBank provides the most comprehensive assembly archive, but quality, completeness, and associated metadata vary enormously across deposited records, and no curated archaeal genome subset is available.

The Genome Taxonomy Database (GTDB) [12] is an ongoing census of bacterial and archaeal diversity through a across the archaeal domain phylogenetically consistent, rank normalized, genome-based taxonomy. It offers archaeal genomes of selective taxonomic representatives under a reclassified, phylogenetically coherent taxonomy. This is an extremely valuable resource, but GTDB serves prim as a taxonomy framework reference rather than a downloadable, quality-filtered, environmental data linked archaeal genomes collection. Besides, GTDB reclassified taxonomy diverges systematically from the latest NCBI taxonomy, creating incompatibilities with the NCBI accessions, BioSample metadata, and taxon identifiers used by many widely deployed bioinformatic tools.

The Integrated Microbial Genomes and Microbiomes system (IMG/M) [13], maintained by the Joint Genome Institute (JGI), contains a large collection of archaeal MAGs with rich environmental context, but is not publicly downloadable as a precompiled genome set with standardized metadata, limiting its utility in large-scale command-line workflows. Collectively, no existing resource provides a dedicated, quality-filtered, environmentally annotated archaeal genome reference set that is directly downloadable and compatible with standard bioinformatic pipelines.

Generally, these challenges underscore the **need** for a robust, quality-controlled archaeal genome resource with broad taxonomic and environmental representation. Such resources are essential to support reliable comparativ and functional of life.

Here we introduce ArchaeaHQ, a quality-controlled, curated reference database of 21,644 archaeal genomes compiled from 35,993 assemblies retrieved from NCBI containing all four archaeal kingdom: Methanobacteriati (Euryarchaeota), Thermoproteati (TACK), Nanobdellati (DPANN), and Promethearchaeati (Asgard). ArchaeaHQ is designed for direct use in metagenomic classification, biogeochemical and ecological context studies, comparative genomics, and the development of archaeal-specific computational tools.

## Results and Discussion

### ArchaeaHQ: A High-Quality Archaeal genome Database

Publicly available genome repositories such as NCBI GenBank are expanding rapidly in the number of archaeal assemblies deposited, but the quality, completeness, and associated environmental metadata of these assemblies vary enormously. No dedicated, systematically curated archaeal genome reference set has previously been available to support environmental and ecological research applications. The initial step of the ArchaeaHQ development was the recovery of archaeal genome assemblies deposited in NCBI as of February 2025, followed by a multi-step curation pipeline (Fig. 1a) that applied standardized quality thresholds to remove species redundancy, low-quality assemblies, and retain genome representatives of the full taxonomic and environmental spectrum of the known archaeal domains.

**Figure 1.**
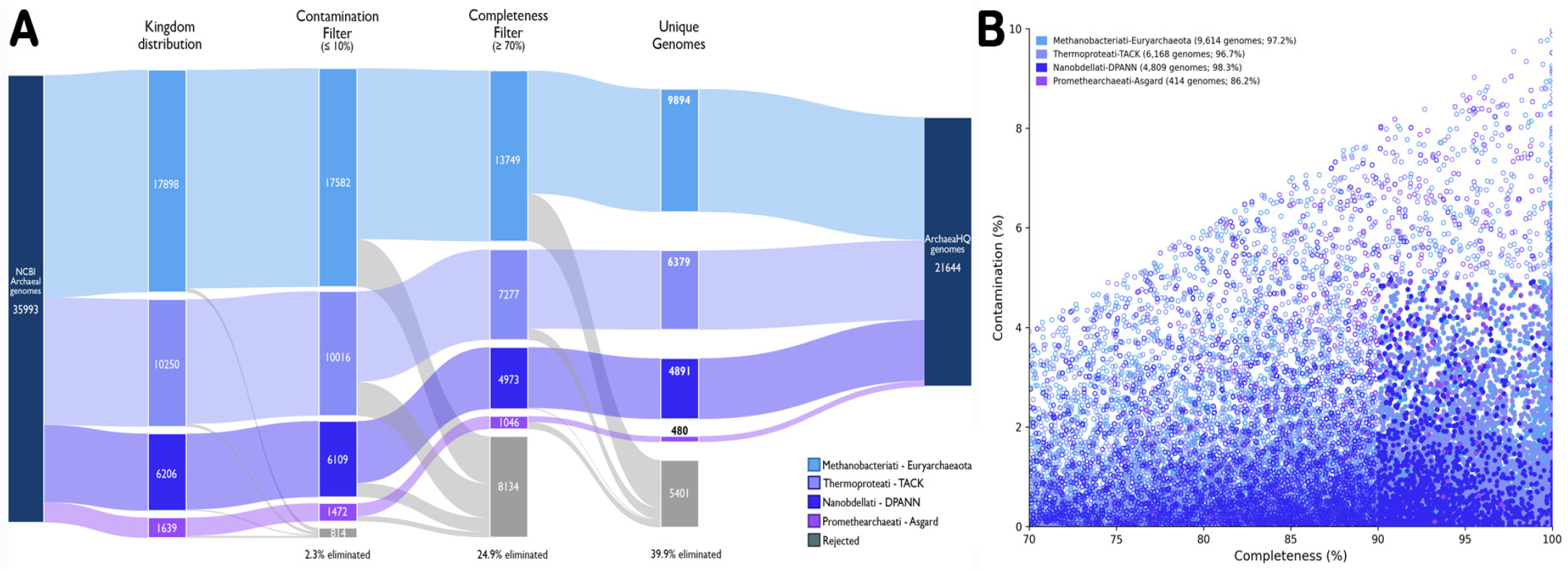
ArchaeaHQ captured a high-quality archaeal genome set after systematic curation removed nearly 40% of NCBI archaeal assemblies. (a) Sankey diagram showing the filtering and curation workflow used to select genomes included in ArchaeaHQ. (b) Completeness and contamination distribution of the curated ArchaeaHQ genomes.. Bars in A and points in B are color-coded by archaeal kingdom: light blue, Methanobacteriati-Euryarchaeota; light purple, Thermoproteati-TACK; dark purple, Promethearchaeati-Asgard; dark blue, Nanobdellati-DPANN. The final curated metadata for all 21,644 archaeal genomes included in ArchaeaHQ are provided in Supplementary Table 1.

The final database comprised 21,644 genomes, a 39.8% reduction from the 35,993 assemblies initially recovered from NCBI. The database includes both cultivated isolate genomes (n = 5,445; 25.2%) and MAGs (n = 16,199; 74.8%), highlighting the fact that the vast majority of archaeal diversity remains uncultivated and it must be accessed through culture-independent approaches.

The initial NCBI dataset contains assemblies from 29,378 unique species-level taxa from all four archaeal kingdoms, confirming that the raw archive captures substantial archaeal species diversity. After excluding identical genomes, ArchaeaHQ comprised the following high quality, unique genome assemblies: Euryarchaeaota (n = 9,894), TACK (n = 6,379), DPANN (n = 4,891), and Asgard (n = 480).

### Genome Quality Metrics

The quality metrics of the retained genomes in ArchaeaHQ reflect the stringency of the pipeline curation (Fig. 1a) and provide a robust foundation for downstream analyses that requires a reliable assembly completeness and low levels of sequence contamination. Completeness across the 21,644 genomes ranges from 70% (the minimum threshold) to 100%, with a mean of 87.7% and a median of 88.4%. A total of 9,558 genomes (44.2%) reach ≥90% completeness, meeting the completeness sub-criterion of the Minimum Information about a Metagenome-Assembled Genome (MIMAG) high-quality standard [14]. Contamination is correspondingly low: values range from 0% to 10% (the maximum threshold), with a mean of 1.52% and a median of 0.78%, and 20,153 genomes (93.1%) exhibit ≤5% contamination. A total of 97.2% of Euryarchaeaota, 96.7% of TACK, 98.3% of DPANN, and 86.2% of Asgard genomes present in ArchaeaHQ have a completeness higher than 90% and contamination lower than 5% (Fig. 1b).

### Environmental Diversity Captured in ArchaeaHQ

The in-depth curation of environmental context of the genomes is what sets ArchaeaHQ apart from other archaeal genome resources, making it directly applicable to ecological and biogeochemical research (Supplementary Table 1). Of the 21,644 genomes in the database, 16,199 (74.8%) are MAGs recovered from environmental metagenomes, while 5,445 (25.2%) are genomes of cultivated isolates. This predominance of MAGs in the dataset reflects the current state of archaeal genomics where cultivation-independent sequencing has been the primary driver of novelty in the archaeal community, particularly for lineages belonging to the DPANN and Asgard kingdoms that have few cultured representatives that require extensive time and resources for **their** cultivation [15 16. the inclusion of both MAGs and isolate-derived genomes in ArchaeaHQ ensures that the database captures both the genomic robustness present in isolated sequencing and the ecological context provided by metagenomic resources.

The environmental metadata provided by ArchaeaHQ (Supplementary Table 2) reveals strong representation of extreme and geochemically active environments. Terrestrial hot springs represent the largest environmental source (n = 3,792 genomes), followed by terrestrial soil (n = 2,282), marine water column (n = 2,128) and deep-sea hydrothermal vent (n = 1,883) (Fig. 2). The dominance of geochemically extreme habitats is consistent with the historical report of high relative abundance of archaeal lineages in these extreme environments, making the recovery of these MAGs less complex than in environments where the bacterial population is dominance is extremely high [17, 18]. The high number of high-quality genomes from extreme environments highlight the value of ArchaeaHQ for studying archaeal contributions to biogeochemical cycling. TACK archaea are particularly well represented in terrestrial hot spring (39.8%), terrestrial soil (16.4%) and hydrothermal vent (12%) assemblies, while Euryarchaeota can be found in a broad array of environments, most of its assemblies are originated from the marine water column (16.3%), terrestrial soil (10%) and animal-host associated samples (10%). DPANN genomes are mainly recovered from subsurface environments (26.9%), acid mine drainage (16.9%), and terrestrial hot springs (14.1%), while Asgard genomes are predominantly recovered from marine sediment (34.8%), mangroves (21.5%), and cold seep systems (11.5%) (Supplementary Table 2).

**Figure 2.**
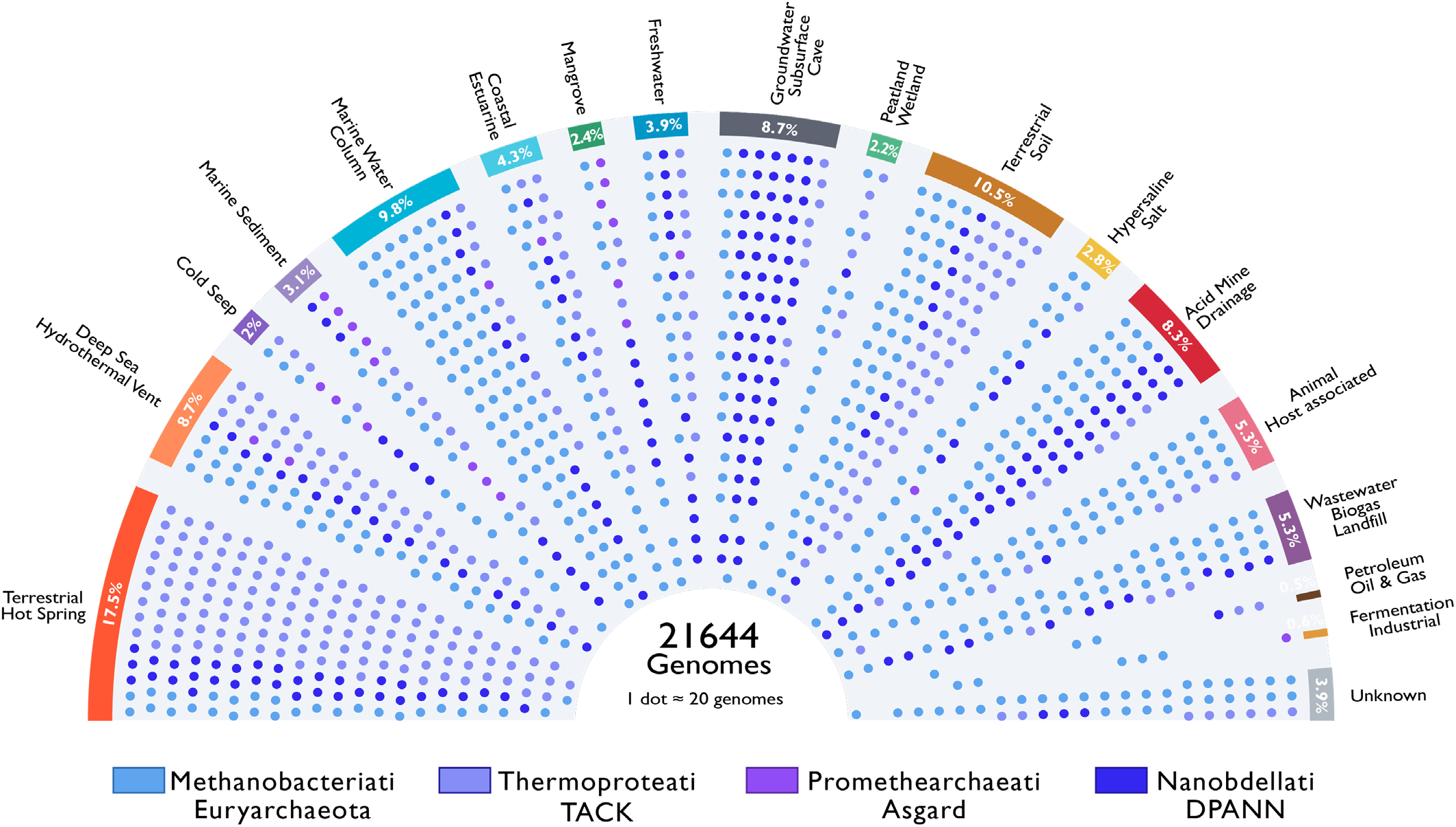
ArchaeaHQ genomes are strongly represented by hot springs, soils, marine waters, and hydrothermal vents. The values on the bars below each environment represent the percentage of all archaeal genomes within that environmental category. Poins under the area covering each environment represent roughly 20 genomes of the corresponding archaeal kingdom and are color-coded by archaeal kingdom: light blue, Methanobacteriati-Euryarchaeota; light purple, Thermoproteati-TACK; dark purple, Promethearchaeati-Asgard; dark blue, Nanobdellati-DPANN. The raw data used to produce this figure can be found in the Supplementary Table 3.

Beyond genomes recovered from extreme environments, ArchaeaHQ also contains several genomes recovered from soil (n = 2,282), marine (n = 2,128 from water column; n = 675 from marine sediment), freshwater (n = 844), and coastal/ estuarine (n = 924) environments which provides a robust foundation to explore biogeochemical cycle dynamics. Archaea in these classic biomes play major roles in carbon, nitrogen and sulfur cycling. ArchaeaHQ leverages high-quality genomes of key biogeochemical players to enable quantitative genomic analyses, metabolic pathway reconstruction, and a de understanding of archaeal functional evolution.

Geographic metadata are available for 14,168 genomes (65.4% of the database), enabling spatial analyses of archaeal biogeography and facilitating the integration of ArchaeaHQ genomes with environmental contextual data in macroecological and comparative genomic studies (Fig. 3). The geographic annotation of ArchaeaHQ assemblies is a direct outcome of the curation workflow, which verified and retained biosample-level metadata from NCBI records, and represents a practical advantage over unfiltered NCBI downloads where metadata completeness is highly variable. To accompany the environmental metadata, the 16S rRNA gene sequence of all genomes in ArchaeaHQ are available in Supplementary table 4 as a resource to facilitate the archaea diversity comparison of specific environments and uses samples.

**Figure 3.**
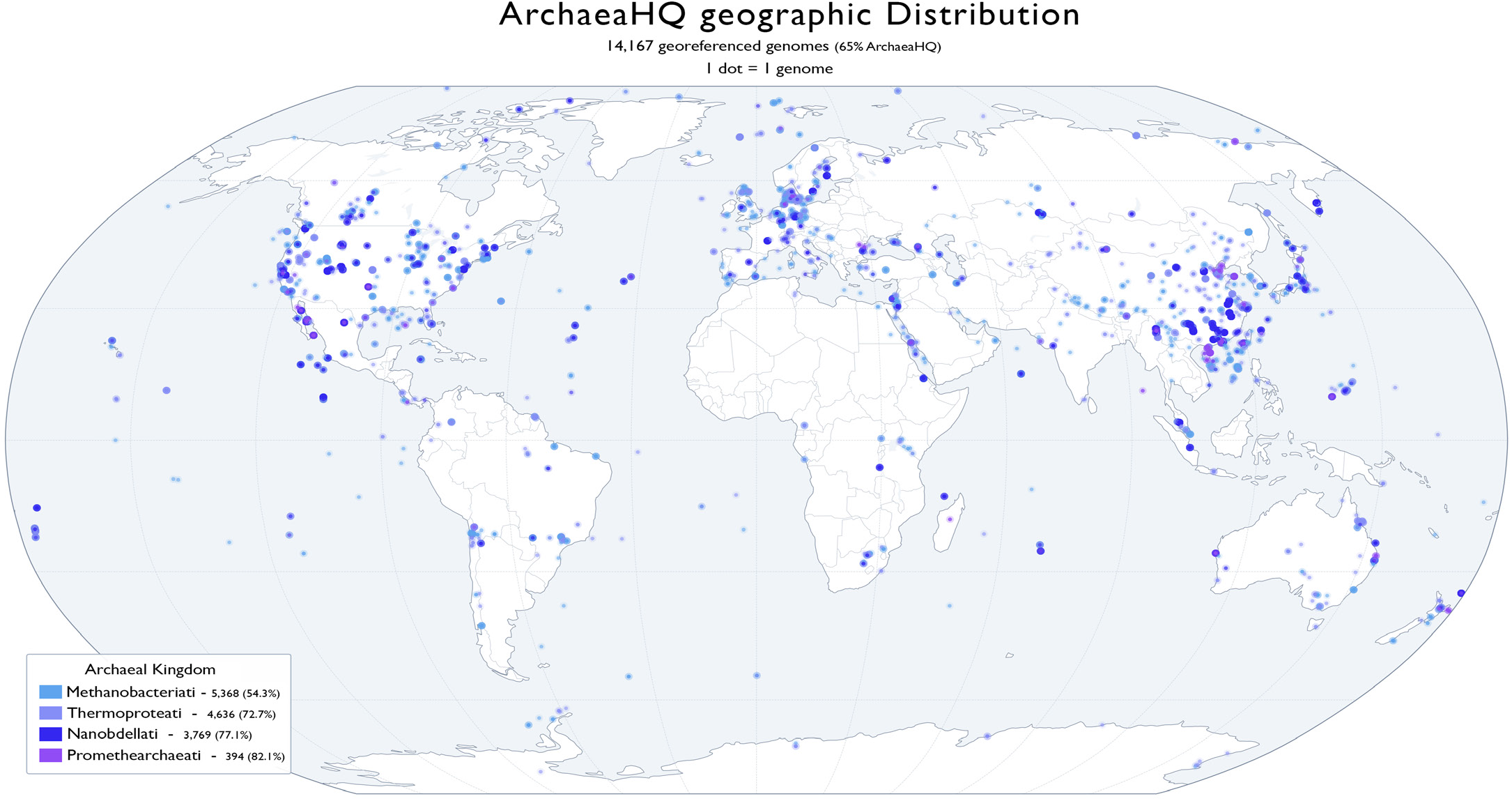
Geographic distribution of the sampling location associated with 14,167 of archaeal genomes present in ArchaeaHQ (65%). Each dot represent one genome, with regions where multiple genomes were isolated from present a growing halo proportional to the amount of genomes isolated from that coordinate. Points are color-coded by archaeal kingdom: light blue, Methanobacteriati-Euryarchaeota; light purple, Thermoproteati-TACK; dark purple, Promethearchaeati-Asgard; dark blue, Nanobdellati-DPANN All the geographic information used in this graph can be found in Suplementary Table 1.

Altogether, the global environmental diversity combined with curated quality control and taxonomic information, positions ArchaeaHQ as a valuable resource for archaeal genomics, with direct applications in metagenomic classification, biogeochemical functional gene surveys, comparative functional annotation, and the development of archaeal-specific bioinformatics tools, including machine learning models for gene prediction and functional annotation.

### Relationship with Existing Archaeal Genomic Resources

Current archaeal genome resources provide valuable but incomplete solutions for large-scale comparative and functional analyses. GTDB (Release 09-RS220) offers a phylogenetically consistent framework with representative genomes but is primarily designed for taxonomy rather than as a downloadable, analysis-ready dataset, and does not prioritize standardized environmental information or compatibility with NCBI-based workflows. The IMG/M contains extensive archaeal genome collections with rich contextual data, yet lacks a unified, quality-filtered, and an easily accessible reference set suitable for direct use in scalable bioinformatic pipelines.

ArchaeaHQ addresses these limitations by providing a curated, quality-controlled, and fully downloadable archaeal genome database ready to be used. It maintains compatibility with NCBI taxonomy and publicly available accessions (e.g. Bioproject, Biosample) (Supplementary Table 1), applies consistent quality thresholds while preserving diversity from underrepresented lineages such as DPANN and Asgard groups, and integrates environmental information as a core feature. By combining accessibility and standardization, ArchaeaHQ fills a critical gap between taxonomy-focused resources and fragmented genome repositories, enabling a reliable method for the incorporation of archaea in future comparative genomics studies and the development of computational biology tools.

## Methods

### Genome retrieval and dataset compilation

All archaeal genome assemblies deposited in NCBI GenBank were retrieved in February 2025 using the NCBI Datasets command-line interface (v.15.19.1) [19] with the `datasets download genome taxon` command. Assemblies were downloaded independently for each of the four recognized archaeal kingdoms: Methanobacteriati (Euryarchaeota; taxon ID 3366610), Thermoproteati (TACK; taxon ID 1783275), Nanobdellati (DPANN; taxon ID 1783276), and Promethearchaeati (Asgard; taxon ID 1935183), yielding a combined dataset of 35,993 genome assemblies. Genome sequences (FASTA format) and associated BioSample metadata, including NCBI assembly accession, BioProject, BioSample identifier, NCBI taxonomic assignment, isolation source, metagenome source label, broad and local environment descriptors (ENVO terms), and geographic coordinates (geo_loc_name, lat/lon), were extracted for each assembly using custom scripts from the genome-metadata-toolkit (https://github.com/Pedrolleao/genome-metadata-toolkit).

### Genome quality assessment

Genome completeness and contamination were estimated for all assemblies using CheckM2 v1.0.1 [20] with default parameters and using the general archaeal marker gene set. Assemblies were retained for downstream analysis only if they simultaneously presented both completeness ≥ 70% and contamination ≤ 10%. These thresholds represent a permissive but meaningful quality gate that captures the range of archaeal genomic diversity while excluding highly fragmented or heavily contaminated assemblies. The 70% completeness minimum sits between high and medium-quality threshold of the MIMAG standard [14], while contamination ≤ 10% is more permissive allowing us to capture a broad diversity of archaeal genomes while eliminating heavily contaminated assemblies.

### Redundancy removal and representative genome selection

Inter-genome comparisons and species-level clustering were performed using dRep v3.6.2 [21]. Primary clustering was performed using Mash distance sketches; within each primary cluster, pairwise average nucleotide identity (ANI) and reciprocal alignment coverage were computed using NUCmer v.4.0.1 [22]. Two tiers of genomic redundancy were resolved using a Union-Find (disjoint-set) algorithm to maintain consistent clustering relationships:

i. Identity-level redundancy (duplicate submissions): genome pairs with ANI ≥ 0.999 and reciprocal alignment coverage ≥ 0.99 were grouped as identical. Because the same assembly is frequently submitted to both NCBI GenBank (GCA\_ prefix) and NCBI RefSeq (GCF\_ prefix), and because minor differences in contig headers or sequence-level edits between the two versions can reduce measured alignment coverage below the threshold even when sequence identity is exact, any pair sharing the same numeric accession identifier in GenBank and Refseq was additionally treated as identical, regardless of measured pairwise coverage. When selecting the preferred representative to keep in ArchaeaHQ from an identity group containing a GCA/GCF pair, the most recent RefSeq version was preferred in the absence of a dRep-designated representative, as RefSeq assemblies undergo additional curation by NCBI.
ii. Species-level redundancy: genome pairs with ANI ≥ 0.95 and reciprocal alignment coverage ≥ 0.95 were assigned to the same species-level cluster, consistent with the widely applied prokaryotic species boundary [23]. Within each species-level cluster, the genome selected by dRep as representative based on a scored composite metric incorporating completeness, contamination, and assembly contiguity (contig N50) was retained in ArchaeaHQ. For species clusters lacking a dRep representative (e.g., due to all cluster members being outcompeted by a genome of higher quality in the ANI comparisons), the most recently curated RefSeq version within the cluster was retained. Genomes not grouped with any other assembly during Mash-distance primary clustering (singletons) were retained directly in ArchaeaHQ as unique representatives.

### Geographic metadata curation

Geographic coordinates were extracted directly from the lat\_lon field of NCBI BioSample records associated with each retained assembly. Records were flagged as georeferenced only when a valid, parseable decimal degree coordinate pair was present; entries containing placeholder values (e.g., “0 N 0 E”, missing , or equivalent uninformative strings) were excluded. Geographic coverage was computed per archaeal kingdom to characterize differential metadata availability across the domain archaeal.

## Data Availability

ArchaeaHQ is available at https://doi.org/10.6084/m9.figshare.32266599. All genome assemblies are publicly available through NCBI GenBank under their original accession numbers. The custom scripts used for metadata extraction, environmental classification, and geographic mapping are available at https://github.com/Pedrolleao/genome-metadata-toolkit.

